# Facile Synthesis and Metabolic Incorporation of *m*-DAP Bioisosteres Into Cell Walls of Live Bacteria

**DOI:** 10.1101/2020.04.03.023671

**Authors:** Alexis J. Apostolos, Julia M. Nelson, Marcos M. Pires

## Abstract

Bacterial cell walls contain peptidoglycan (PG), a scaffold that provides proper rigidity to resist lysis from internal osmotic pressure and a barrier to protect cells against external stressors. It consists of repeating sugar units with a linkage to a stem peptide that becomes highly crosslinked by cell wall transpeptidases (TP). Because it is an essential component of the bacterial cell, the PG biosynthetic machinery is often the target of antibiotics. For this reason, cellular probes that advance our understanding of PG biosynthesis and its maintenance can be powerful tools to reveal novel drug targets. While synthetic PG fragments containing L-Lysine in the 3^rd^ position on the stem peptide are easier to access, those with *meso*-diaminopimelic acid (*m*-DAP) pose a severe synthetic challenge. Herein, we describe a solid phase synthetic scheme based on the widely available Fmoc-protected L-Cysteine building block to assemble *meso*-cystine (*m*-CYT), which mimics key structural features of *m*-DAP. To demonstrate proper mimicry of *m*-DAP, cell wall probes were synthesized with *m*-CYT in place of *m*-DAP and evaluated for their metabolic processing in live bacterial cells. We found that *m*-CYT-based cell wall probes were properly processed by TPs in various bacterial species that endogenously contain *m*-DAP in their PG. We anticipate that this strategy, which is based on the use of inexpensive and commercially available building blocks, can be widely adopted to provide greater accessibility of PG mimics for *m*-DAP containing organisms.

## Introduction

Peptidoglycan (PG) is a structurally integral part of the bacterial cell wall that provides cellular rigidity and prevents lysis from osmotic stress. The primary monomeric unit of PG includes a disaccharide of *N*-acetylglucosamine (GlcNAc) and *N*-acetylmuramic acid (MurNAc) sugars, with linkage to a stem peptide, typically a pentamer with the sequence L-Ala-iso-D-Glu-L-Lys-D-Ala-D-Ala or with *meso*-diaminopimelic acid (*m*-DAP) in place of L-Lys at the 3^rd^ position.**Error! Reference source not found.** Nascent PG units are imbedded within the existing PG scaffold by the transglycosylation of the disaccharides and the crosslinking of neighboring stem peptides, carried out by transpeptidase (TP) enzymes.**Error! Reference source not found.**

There are considerable similarities in PG structure across different types of bacteria and, yet, slight structural variations can alter cell wall physiochemical properties and tune its interaction with neighboring cells including host organisms. Today, we know that fragments of PG are released into their surrounding environments as by-products of cell wall metabolism or as a component of signaling systems to modulate the response from a neighboring bacteria or a host organism.^1-3^ A prominent difference in PG primary structure happens on the 3^rd^ position on the stem peptide, which is also the site for PG crosslinking and Braun’s lipoprotein attachment (**Figure 1**).^4, 5^ The difference of a single carboxylic acid between *m*-DAP and L-Lys can play a major role in how PG is sensed. As an example, vegetative *Bacillus subtilis* (*B. subtilis*) cells release PG fragments to communicate the initiation of a synchronized germination event from spores using the sensor protein PrkC.^6, 7^ The specificity in this communication is dictated by the complete lack of PrkC response to L-Lys containing PG fragments, which is nonnative to the *m*-DAP based PG of *B. subtilis*.^8^ Similarly, bacteria release biomacromolecules (e.g., PG) that are collectively known as pathogen-associated molecular patterns (PAMPs), which can be sensed by human immune cells indicating that there may be an infection.^9^ NOD1, from the family of PAMP receptors, is known to be activated by the minimal agonist D-Glu-*m*-DAP (iE-DAP).^10^ It was recently demonstrated that several bacterial species amidate *m*-DAP to evade NOD1 activation.^11, 12^ Together, these examples highlight how alterations to the PG stem peptide structure can regulate sensing and metabolism of PG fragments.

**Figure 1.**
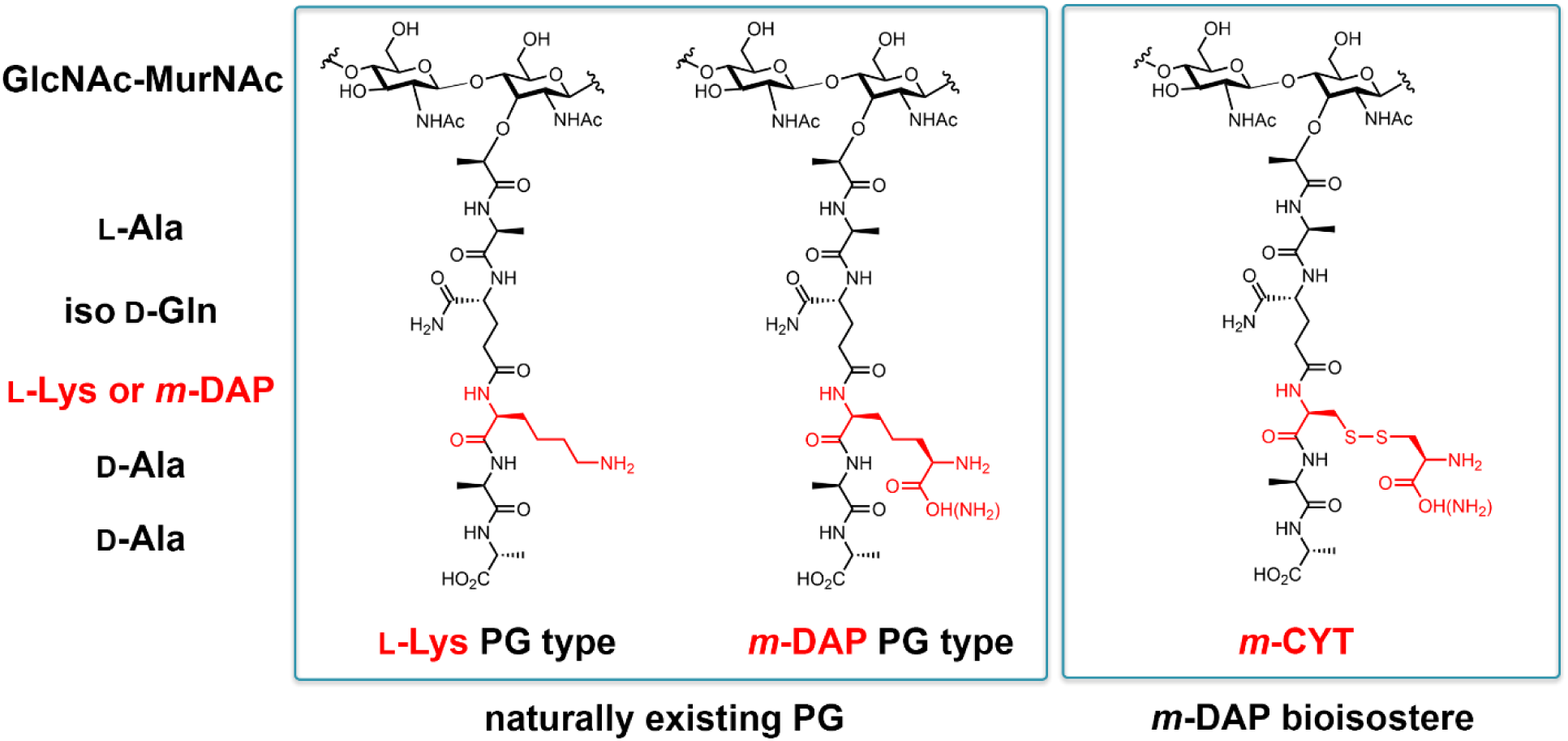
Chemical structure of the basic unit of PG (disaccharide attached to the stem pentapeptide). A major difference between bacterial species occurs at the 3^rd^ position (L-Lys and *m*-DAP). This schematic diagram shows how the bioisostere *m*-CYT mimics key structural features of *m*-DAP.

A large number of pathogenic and commensal bacteria have *m*-DAP based PG. Therefore, it is critically important to be able to synthesize homogeneous *m*-DAP based PG fragments (as opposed to heterogenous isolated PG fragments from live bacteria) to systematically dissect the biological consequences of alterations to the PG stem peptide found in these organisms.^13, 14^ While Fmoc-protected L-Lys building blocks are readily available and highly compatible with standard solid phase peptide chemistry, *m*-DAP is not commercially available and this poses a significant synthetic barrier. The internal symmetry of *m*-DAP coupled with the need to have differentiated, orthogonally protected stereocenters, makes its synthesis and that of its protected counterparts highly challenging.^15-20^ Instead of the difficult-to-synthesize *m*-DAP, an alternative strategy involves the utilization of *m*-DAP analogs that mimic the key components and are functionally similar to *m*-DAP. Examples of a such analogs include *meso*-lanthionine (*m*-LAN)^21-23^ and *meso*-oxa-DAP.^24-27^ While both of these analogs recapitulate aspects of *m*-DAP as demonstrated by their processing by enzymes that recognize *m*-DAP residues, their synthesis still involves several solution phase chemistry steps and they are not readily adaptable to Fmoc-based solid phase chemistry. We envisioned that we could use widely available Fmoc-protected L-Cys as a precursor to the synthesis of *meso*-cystine (*m*-CYT) analogs (**Figure 1**). *m*-CYT preserves critical structural features at the terminal end of the sidechain and this building block can easily be diversified using commercially available amino acids to build homogenous PG fragments and their naturally occurring derivatives. We leveraged our synthetic strategy to access variants of *m*-DAP bioisosteres and showed that naturally existing modifications to the PG structure can alter their metabolism and processing by cell wall biosynthetic machinery.

At first, we set out to establish two potential routes to build *m*-CYT containing PG fragments: one that involved a post-cleavage modification with the crude material and the other that involved the installation of the sidechain on solid support (**Scheme 1**). Synthesis of the pentapeptide with an *N*-terminal MurNAc was carried out using commercially available building blocks. In route *A*, Fmoc-L-Cys(Trt)-OH was coupled into the 3^rd^ position of the pentapeptide using conventional peptide coupling conditions. Following the concomitant global deprotection and release from the solid support, the crude material was treated with D-cystine to promote the installation of D-Cys onto the free sulfhydryl of the L-Cys containing peptide *via* disulfide exchange to generate *m*-CYT. Alternatively in route *B*, a L-Cys residue protected with a acetamidomethyl (ACM) sidechain was installed on the 3^rd^ position of the pentapeptide.^28^ Once all coupling reactions were complete, the ACM moiety was selectively converted to *S*-carbomethoxysulfenyl (Scm) on resin and converted to the asymmetric disulfide by treatment with D-Cys. For both routes, the peptides are subjected to RP-HPLC purification and characterized using HR-MS. We found that both routes *A* and *B* yielded satisfactory levels of the final product (**Figure 2** or convert Scheme 2 to Figure 2 and add panels of analytical HPLC and MS?). The lack of cysteine residues in naturally existing PG will ensure the chemoselective modification of L-Cys in synthesizing *m*-CYT containing PG fragments.

**Figure 2.**
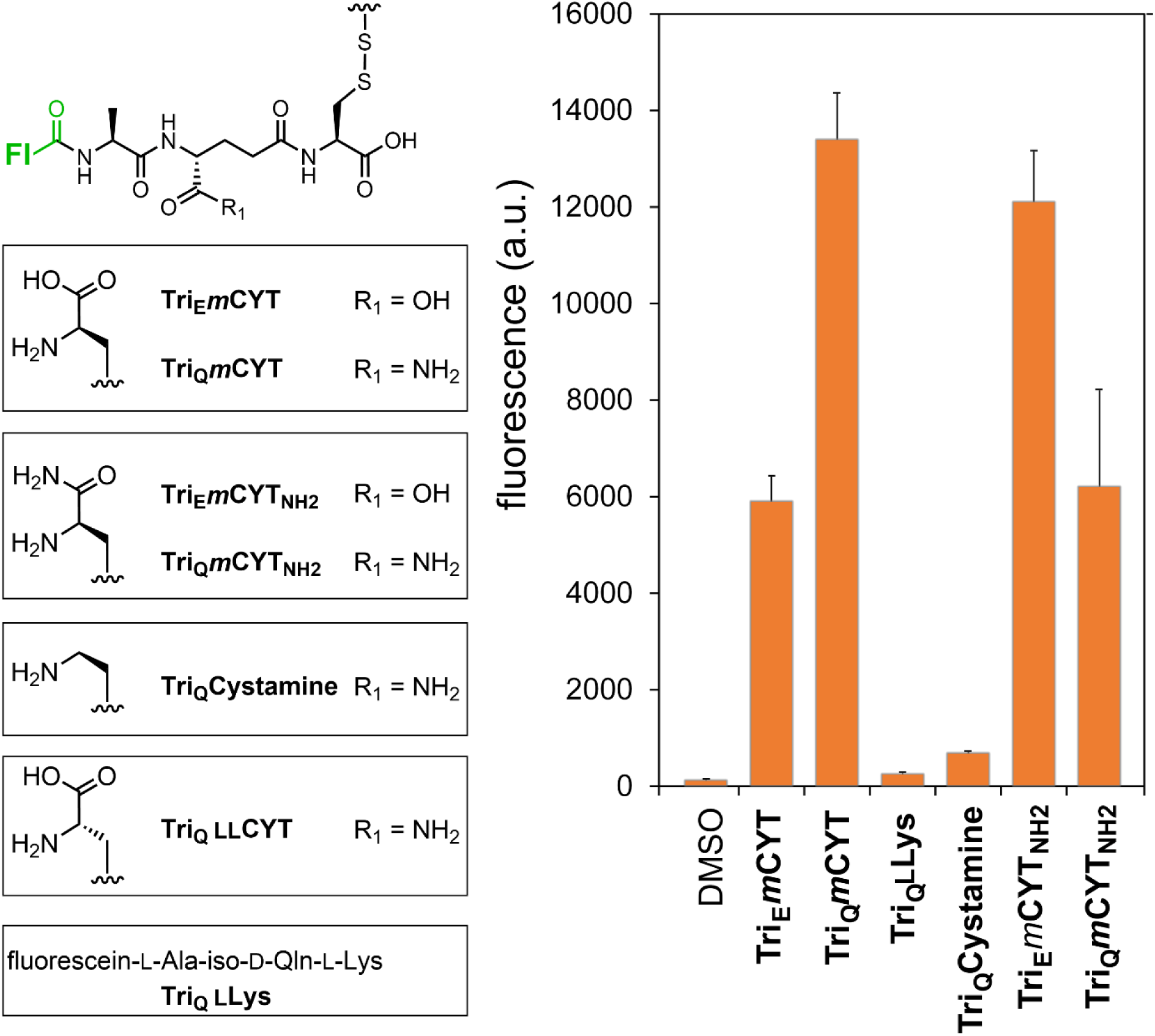
Chemical structures for tripeptide-based probes. Fl = carboxyfluorescein. Flow cytometry analysis of *M. smegmatis* treated overnight with 100 μM of tri-peptide probes. Data are represented as mean + SD (n = 3).

With synthetic strategies in place, we turned our attention to investigating how *m*-CYT would get recognized and processed metabolically by live bacterial cells. We anticipated that we could leverage the mimicry of *m*-DAP by *m*-CYT to study PG crosslinking, which is an essential step in the biosynthesis of properly structured bacterial cell walls. PG crosslinking is mainly carried out by Penicillin Binding Proteins transpeptidases (D,D-TP) and L,D-transpeptidases (Ldts) to generate distinct connectivity patterns (**Scheme 2A**). The importance of these two TP classes of enzymes is illustrated by the general lack of sensitivity of Ldts against most β-lactam antibiotics (e.g., penicillins and cephalosporins). A large number of clinically relevant antibiotics interfere with PG crosslinking and, therefore, probes that track cell wall crosslinking dynamics can potentially reveal novel targets for drug discovery. Single D-amino acid PG probes have now become essential tools to image PG biosynthesis.^29-34^ Yet, single D-amino acid PG probes cannot interrogate how the stem peptide structure (or its many chemical modifications) can control PG crosslinking or processing. We^35, 36^, and others^37-40^, have recently demonstrated that fluorescently tagged structural mimics of nascent TP substrates can be crosslinked into the growing PG scaffold of live cells, thus hijacking the cell wall machinery (**Scheme 2B**). Indeed, we have shown that penta-, tetra-, and tri-peptides that are analogs of the PG stem peptide are accepted by TPs to varying extents.^35, 36^ However, to this point we had been unable to investigate the role of *m*-DAP (and its modifications) because of the challenge in building *m*-DAP containing peptides.

We first built a series of *m*-CYT PG probes using the synthetic routes we developed *via* Fmoc-based solid phase peptide chemistry. Tripeptides were chosen to establish the recognition of the sidechain since tripeptides cannot act as acyl-donors in the TP reaction (they lack the D-Ala necessary for the acyl-donor step) and, instead, operate solely as acyl-acceptors. In this configuration, the amino group on the *m*-CYT tripeptide acts as the nucleophile in the PG crosslinking reaction (**Scheme 2B**). Fluorescein was conjugated to the *N*-terminus to track incorporation into the PG *via* TPs and fluorescence can be readily quantified using flow cytometry (**Scheme 2C**).

We first chose to evaluate the metabolic labeling of *Mycobacterium smegmatis* (*M. smegmatis*), whose PG biosynthesis tracks similarly with the pathogenic *Mycobacterium tuberculosis*, the causative agent of Tuberculosis (TB).^41-43^ TB continues to impose a tremendous health burden across the globe and the unique composition of the *Mycobacterium* cell wall provides an opportunity for the elucidation of novel drug targets. For this set of experiments, *M. smegmatis* growing in early lag-phase were treated with exogenously supplied probes and incubated overnight. Supplementation of *M. smegmatis* cells with **Tri_E_*m*CYT** (a tripeptide probe that contains iso-D-Glu on the second position and *m*-CYT on the 3^rd^ position) resulted in cellular fluorescence levels that were ∼50-fold higher than control cells, a clear indication that *m*-CYT satisfactorily mimics *m*-DAP and is recognized by the native TP (**Figure 2**). While PG precursors are initially biosynthesized with an iso-D-Glu in the 2^nd^ position of the stem peptide, it is now established that for several Gram-positive organisms, iso-D-Glu is converted to iso-D-Gln.^44-46^ In fact, the gene encoding the tandem enzyme MurT-GatD, responsible for iso-D-Glu amidation, is essential in *Staphylococcus aureus* and *Streptococcus pneumoniae*.^47, 48^ It was demonstrated, *in vitro*, that absence of iso-D-Glu amidation resulted in undetectable levels of PG crosslinking.^44, 45^ Yet, PG crosslinking reactions utilize two PG strands and it was not established if iso-D-Glu amidation was required on both acyl-acceptor and acyl-donor strands. Our group recently showed that iso-D-Glu amidation plays an important role in controlling PG crosslinking in the acyl-donor chain of *M. smegmatis* and *M. tuberculosis*.^36^ The tripeptide based probes lack D-Ala on the 4th and 5th position, which means that they can only be incorporated into the PG as acyl-acceptor strands. Amidation of the second position to iso-D-Glu (**Tri_Q_*m*CYT**) resulted in the doubling of PG incorporation as evident by the increase in cellular fluorescence, which indicates a preference for amidated iso-D-Glu but not an absolute requirement. Our new results indicate that iso-D-Glu amidation in the acyl-acceptor strand may not be as an absolute requirement for PG crosslinking in acyl-acceptor strand of *M. smegmatis*.

The sidechain of the 3^rd^ position (L-Lys or *m*-DAP, **Figure 1**) in naturally existing PG contains the critical amine nucleophile and it is, theoretically, possible that PG strands lacking the carboxylic acid could be competent acyl-strand acceptors in the TP crosslinking reactions. To test the essentiality of the carboxylic acid moiety in *m*-CYT, two control probes were designed and synthesized: **Tri_Q_LLys** and **Tri_Q_Cystamine**. However, our results showed that the carboxylic acid moiety is essential for the acyl-acceptor strand to crosslink into the PG scaffold as the analogous probes lacking the carboxylic acid group led to minimal cellular fluorescence regardless of the linker composition (methylene and disulfide, **Figure 2**). Critically, these results also indicate that the increase in cellular fluorescence is not purely due to disulfide exchange at the cell surface based on the low labeling levels with **Tri_Q_Cystamine**. A major advantage of our strategy is the ability to access derivatives of *m*-DAP that are known to be naturally occurring such as amidation of the carboxylic acid. Currently, there is little known about *m*-DAP amidation in *M. smegmatis* and *M. tuberculosis*, whereas it was previously reported that the PG of *Mycobacterium abscessus* contains about 20 percent of amidated *m*-DAP.^49^ Some plausible biological roles of *m*-DAP amidation have been proposed. In a transposon screen for lysozyme potentiation in a *m*-DAP containing organism, *Rhodococcus erythropolis*, a gene was identified that is related to *m*-DAP amidation.^50^ Later, the same gene was found to play a role in β-lactam antibiotic sensitivity in *Corynebacterium glutamicum* (*C. glutamicum*)^51^ and *B. subtilis*.^52^ The absence of live cell PG probes has hampered our ability to establish the effect of *m*-DAP amidation on PG crosslinking. To gain insight into this open question, we investigated how *m*-DAP amidation impacts TP crosslinking levels by building **Tri_E_*m*CYT_NH2_** and **Tri_Q_*m*CYT_NH2_**. Interestingly, amidation of the *m*-DAP crossbridge in combination with iso-D-Glu (**Tri_E_*m*CYT_NH2_**) led to similar TP-based incorporation as **Tri_Q_*m*CYT**. These results suggest the possibility that *m*-DAP amidation serves as a compensatory modification to the lack of iso-D-Glu amidation. The double amidation in **Tri_Q_*m*CYT_NH2_** led to fluorescence levels equivalent to with **Tri_E_*m*CYT**.

Having demonstrated that tripeptide probes displaying *m*-CYT residues are functional in live bacterial cells, we next performed several additional experiments to further define the mode of probe incorporation into the bacterial PG scaffold. Incorporation of tripeptide-based probes can potentially be processed by two distinct classes of TPs (**Figure 1A**). We attempted to determine if an individual TP class contributed more prominently to the crosslinking of the *m*-CYT probe into the PG scaffold by using antibiotics to interfere with specific TP function. As a point of comparison, we also assessed incorporation of two probes that we had previously disclosed to be D,D-TP specific (**Penta_Q_LLys**) and Ldt-specific (**Tetra_Q_LLys**).^36^ Co-incubation of ampicillin, a β-lactam that does not inhibit Ldts^53^, with **Tri_Q_*m*CYT** led to a slight increase in cellular labeling in a concentration dependent manner consistent with Ldt-mediated PG incorporation and similar to the Ldt-specific **Tetra_Q_LLys** (**Figure 3**). The increase in cellular fluorescence may be interpreted as a response mechanism to the inactivation of parts of the cell wall biosynthetic machinery, a feature that we previously observed with *Enterococcus faecium*^*36*^ and we are exploring further. As expected, ampicillin treatment led to a decrease in cellular labeling with the D,D-TP specific **Penta_Q_LLys**. Treatment with sub-lethal concentrations of meropenem, an inhibitor of both TP classes^54^, resulted in a decrease of **Tri_Q_*m*CYT** labeling. The mechanism of PG incorporation was further interrogated by using Ldt-deletion strains. Whereas deletion of two individual Ldts did not impact cellular labeling, the deletion of the five primary Ldts led to near background cellular fluorescence (**Figure 4**). Together, these results principally established that *m*-CYT is a functional substitute for native *m*-DAP in TP processing in live bacterial cells.

**Figure 3.**
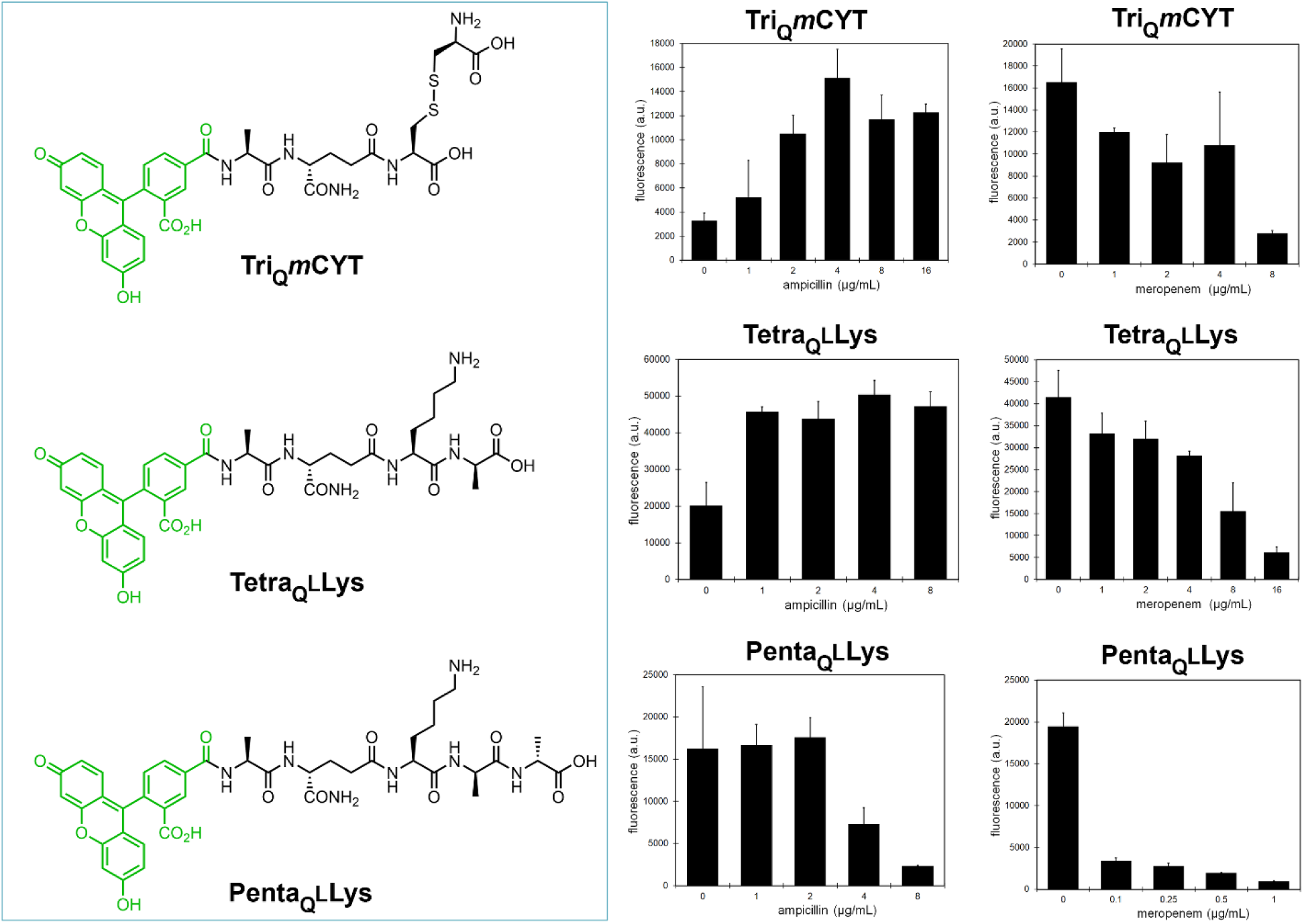
Flow cytometry analysis of *M. smegmatis* treated overnight with 100 μM of each probe and increasing concentrations of ampicillin or meropenem. Data are represented as mean + SD (n=3).

**Figure 4.**
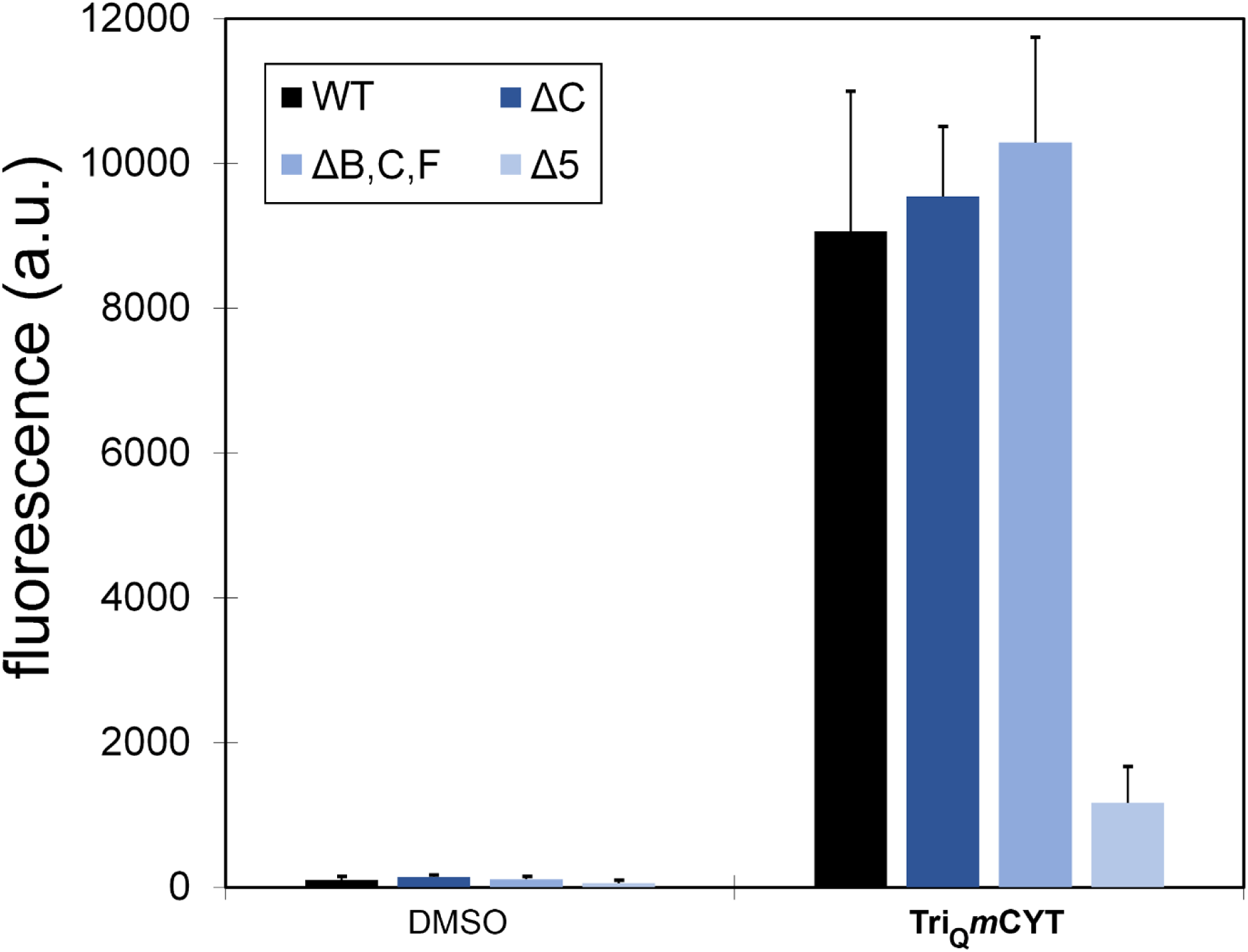
Flow cytometry analysis of *M. smegmatis* (WT) and Ldt knockout mutants untreated or treated overnight with 100 μM **Tri_Q_*m*CYT**. Data are represented as mean + SD (n=3).

Cellular probes can also reveal aspects of bacterial adaptation to human immune responses. In the case of *M. tuberculosis* infections, bacterial cells are phagocytosed by macrophages to promote pathogen destruction in lysosomal compartments. Acidification of the phagosome is a critical component for proper elimination of bacterial pathogens.^55^ In turn, *M. tuberculosis* and *M. smegmatis* have evolved the ability to tolerate acidic environments by reprogramming a series of genetic elements^56^ including some that are associated with PG biosynthesis.^57, 58^ To test the potential for PG remodeling in response to acidic environments, *M. smegmatis* cells were grown in media supplemented with probes at pH 4.0 or pH 7.4 (**Figure 5**). Remarkably, there was a nearly 8-fold increase in fluorescence levels for cells incubated with **Tri_Q_*m*CYT** in acidic media but no induction was observed for **Tri_Q_LLys**. This marked increase in cellular fluorescence could, potentially, suggest two modalities of PG reconfiguration during phagosome-like acidic stress. One possibility is that phagosome-like conditions trigger an upregulation of PG crosslinking that is reflected in higher engraftment of probes, which would be consistent with past evidence of upregulation of PG TPs in *M. smegmatis* challenged with low pH conditions.^59^ Alternatively, higher cellular fluorescence could be a result of downregulation of PG hydrolases, which could result in higher retention of the PG probes following their incorporation.

**Figure 5.**
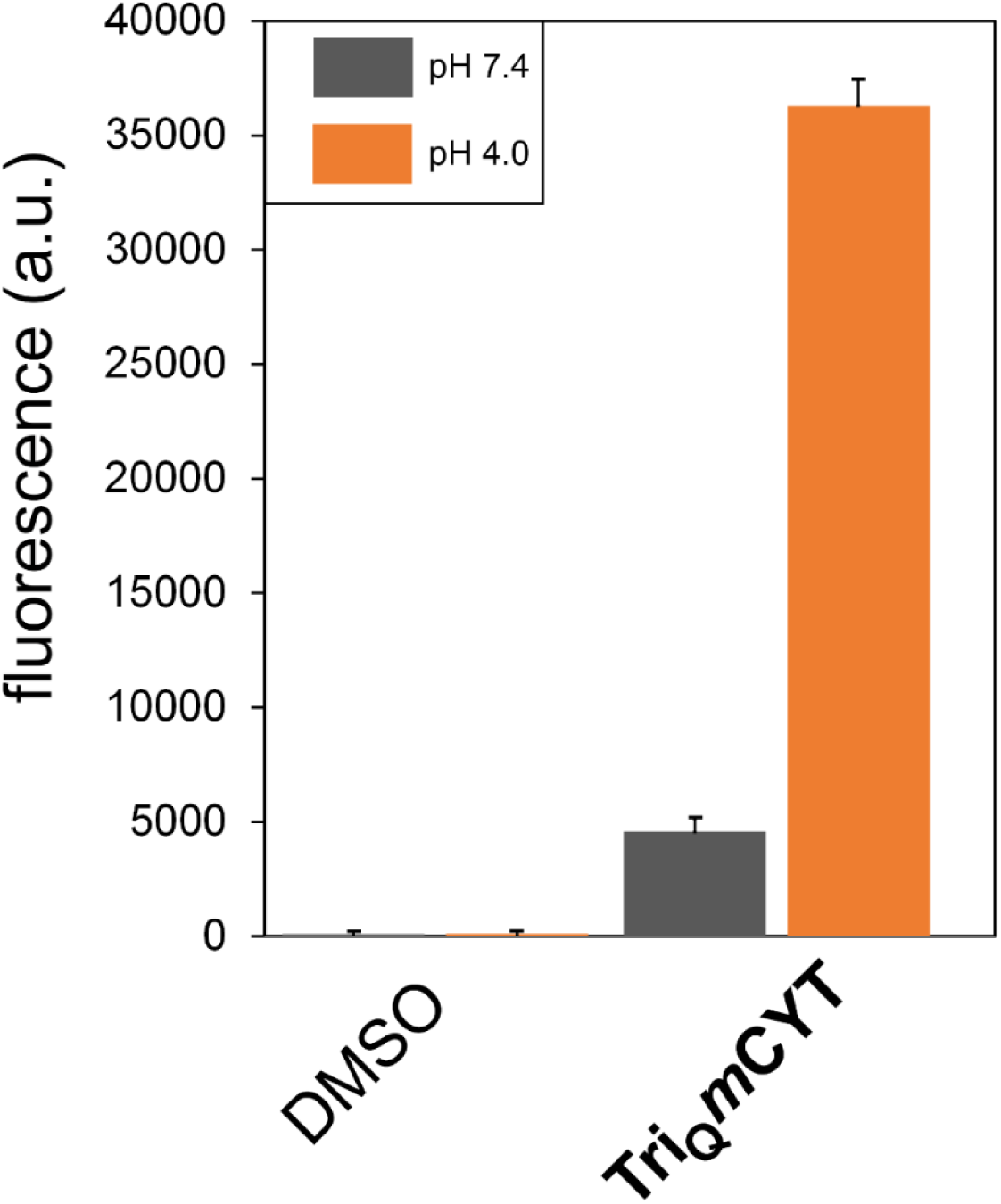
Flow cytometry analysis of *M. smegmatis* treated overnight with 100 μM of **Tri_Q_*m*CYT** at pH 7.4 or pH 4.0 (buffered with 2-[N-morpholino]ethanesulfonic acid [MES]). Data are represented as mean + SD (n=3).

We extended our analysis to probes that cover a wider range of TP substrates under acid stress by analyzing PG incorporation of **Tetra_Q_LLys** and **Penta_Q_LLys** (**Figure 6**). In contrast to **Tri_Q_*m*CYT**, there was no change in PG incorporation of **Tetra_Q_LLys** and there was, in fact, a decrease in incorporation of **Penta_Q_LLys** in acidic conditions. This labeling pattern suggests that induction in PG crosslinking must have an *m*-DAP-like sidechain and it is not indiscriminately elevated at low pH levels. Quite strikingly, re-introducing the carboxylic acid group within **Tetra_Q_*m*CYT** and **Penta_Q_*m*CYT** led to a return in the induction of PG incorporation at low pH levels. The requirement for the carboxylic acid group on the sidechain of *m*-DAP for proper adaptation to acidic environments has been determined for at least one protein, ArfA.^60^ A domain within ArfA, which is proposed to function as a OmpA-like porin, responds to low pH *via* a conformational change to associate with the *m*-DAP sidechain within PG but not L-Lys PG type. With a synthetic route to build *m*-DAP-like probes, in the future we will be uniquely positioned to investigate how our probes containing *m*-CYT may function to report on cell wall remodeling in *Mycobacteria* during phagocytosis.

**Figure 6.**
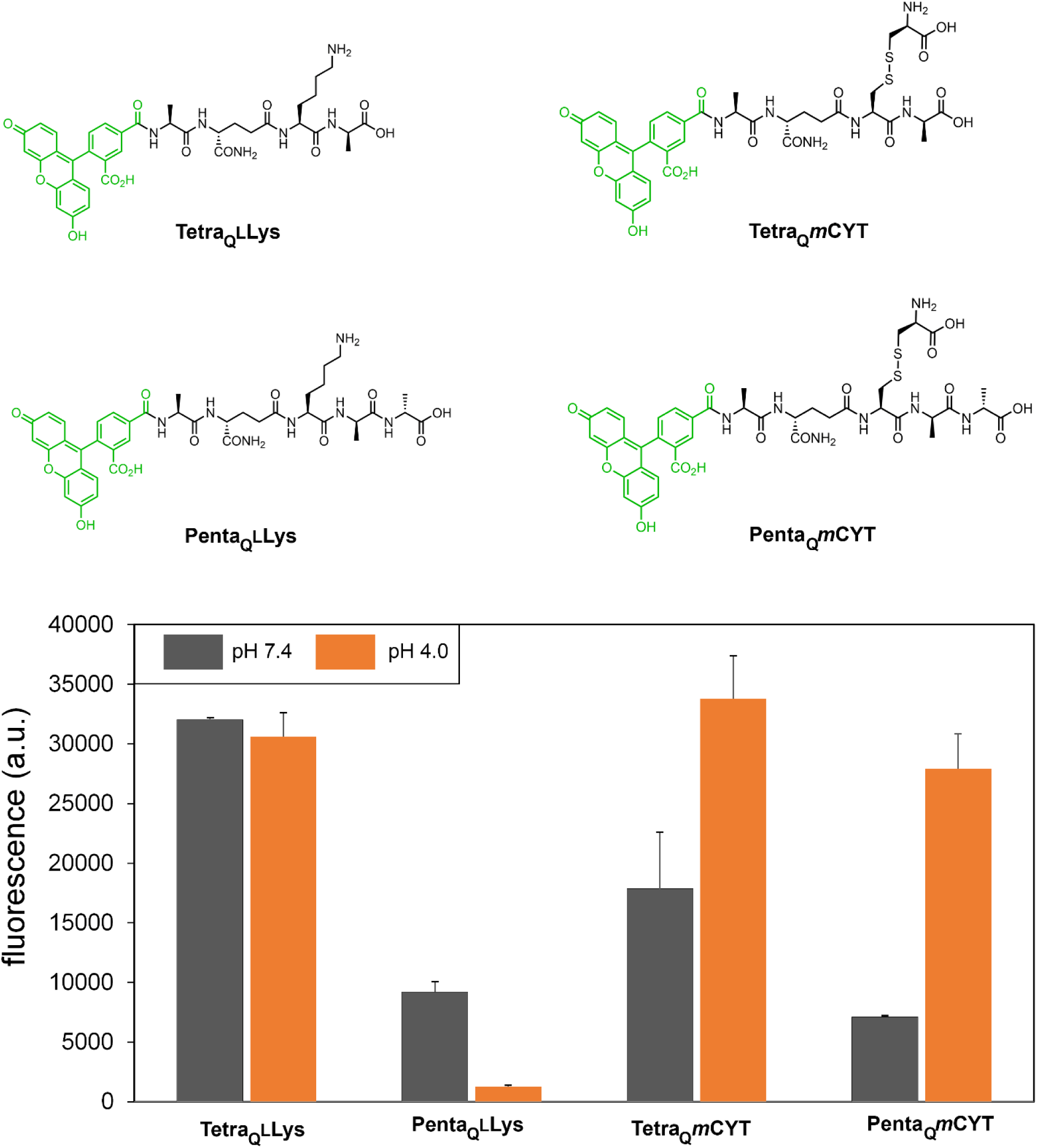
Flow cytometry analysis of *M. smegmatis* treated overnight with 100 μM of each probe at pH 7.4 or pH 4.0 (buffered with 2-[N-morpholino]ethanesulfonic acid [MES]). Data are represented as mean + SD (n=3).

Bacteria use modifications to the primary sequence of the PG stem peptide as an adaptation mechanism that is, likely, specific to its needs and niche.^6, 61^ As such, it is possible that amidation of D-iGlu and *m*-DAP may not have the same consequence in all bacteria with *m*-DAP based PG. We became interested in leveraging our probes to investigate naturally occurring PG modifications in the probiotic organism *Lactobacillus plantarum* (*L. plantarum*).^62^ Despite our lack of understanding of all the ways that probiotics modulate host immune responses, it is clear that PG fragments play an integral part *via* the modulation of NOD receptors.^63, 64^ Chemical modifications to the PG stem peptide can, in turn, alter the level of activation by PG sensors as is evidenced by the muted response by NOD1 receptors when treated with PG fragments containing amidated D-iGlu^65^ and *m*-DAP.^66^ In fact, deletion of the gene responsible for *m*-DAP amidation in *L. plantarum* results in impaired cell growth and filamentation^67^ and it is lethal in *B. subtilis*.^*52*^ Incorporation of tripeptide based probes into the PG of *L. plantarum* was tested by supplementing the probes during cell culture (**Figure 7**). Similar to *M. smegmatis*, the carboxylic acid within the sidechain in the 3^rd^ position of the stem peptide was critical for high cellular labeling levels. In contrast to *M. smegmatis*, there is less pronounced selectivity for D-iGlu amidation and, instead, *m*-DAP amidation alone is sufficient to yield high levels of PG incorporation (**Figure 7**). There was no induction of PG incorporation at acidic pH conditions with *m*-CYT probes, suggesting that the increase in PG labeling in acidic conditions is specific to *Mycobacteria* (**Figure 8**). Interestingly, the supplementation of **Tri_Q_*m*CYT_NH2_** in *L. plantarum* led to the highest labeling levels our laboratory has ever observed with any cell wall probe. The high cell surface labeling levels observed could prove to be helpful in grafting epitopes (in place of fluorescein) on the surface of *L. plantarum* to promote specific adhesion to abiotic surfaces or targeted binding to pathogenic mammalian cells.

**Figure 7.**
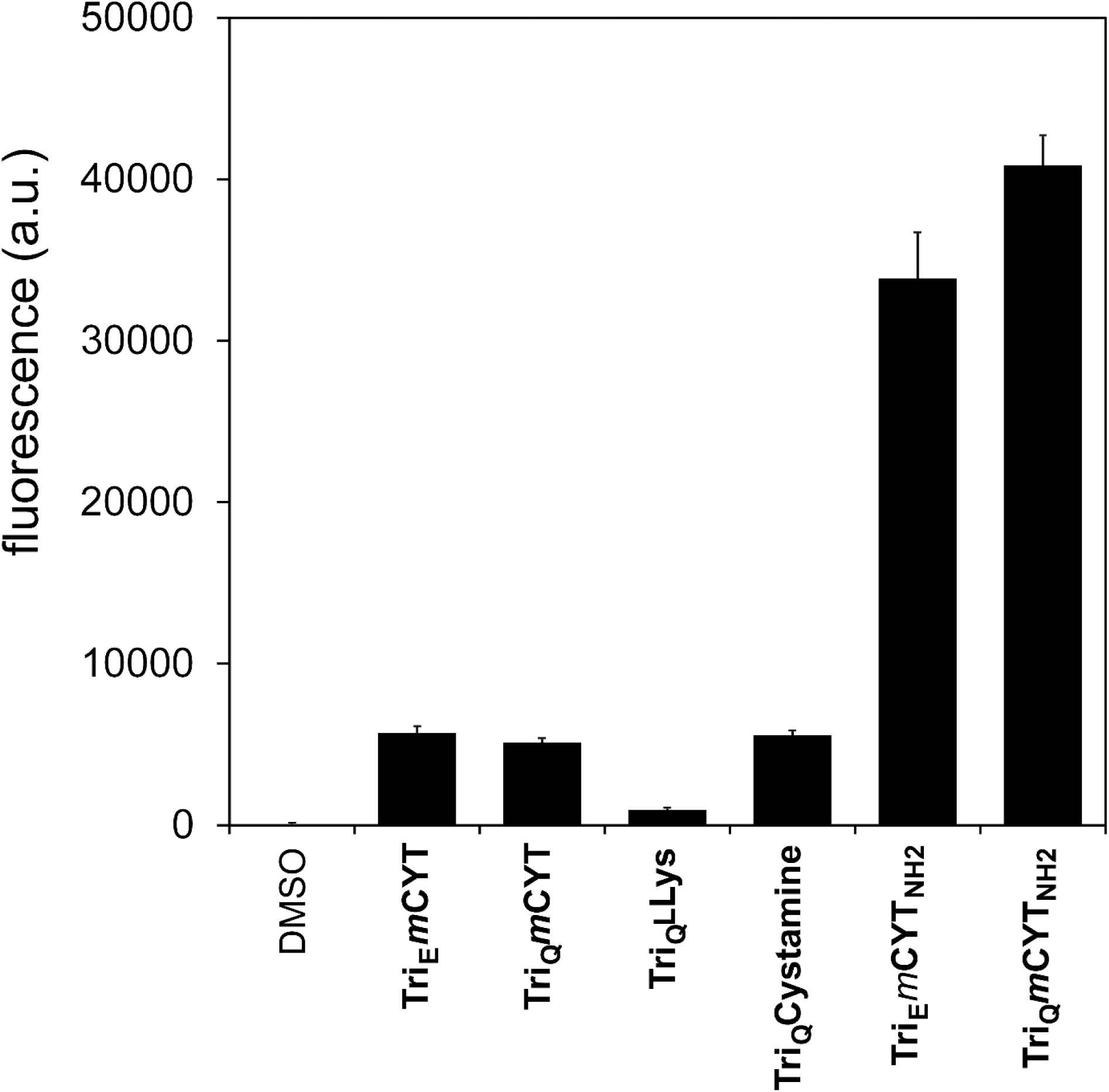
Flow cytometry analysis of *L. plantarum* treated overnight with 100 μM of each probe. Data are represented as mean + SD (n=3).

**Figure 8.**
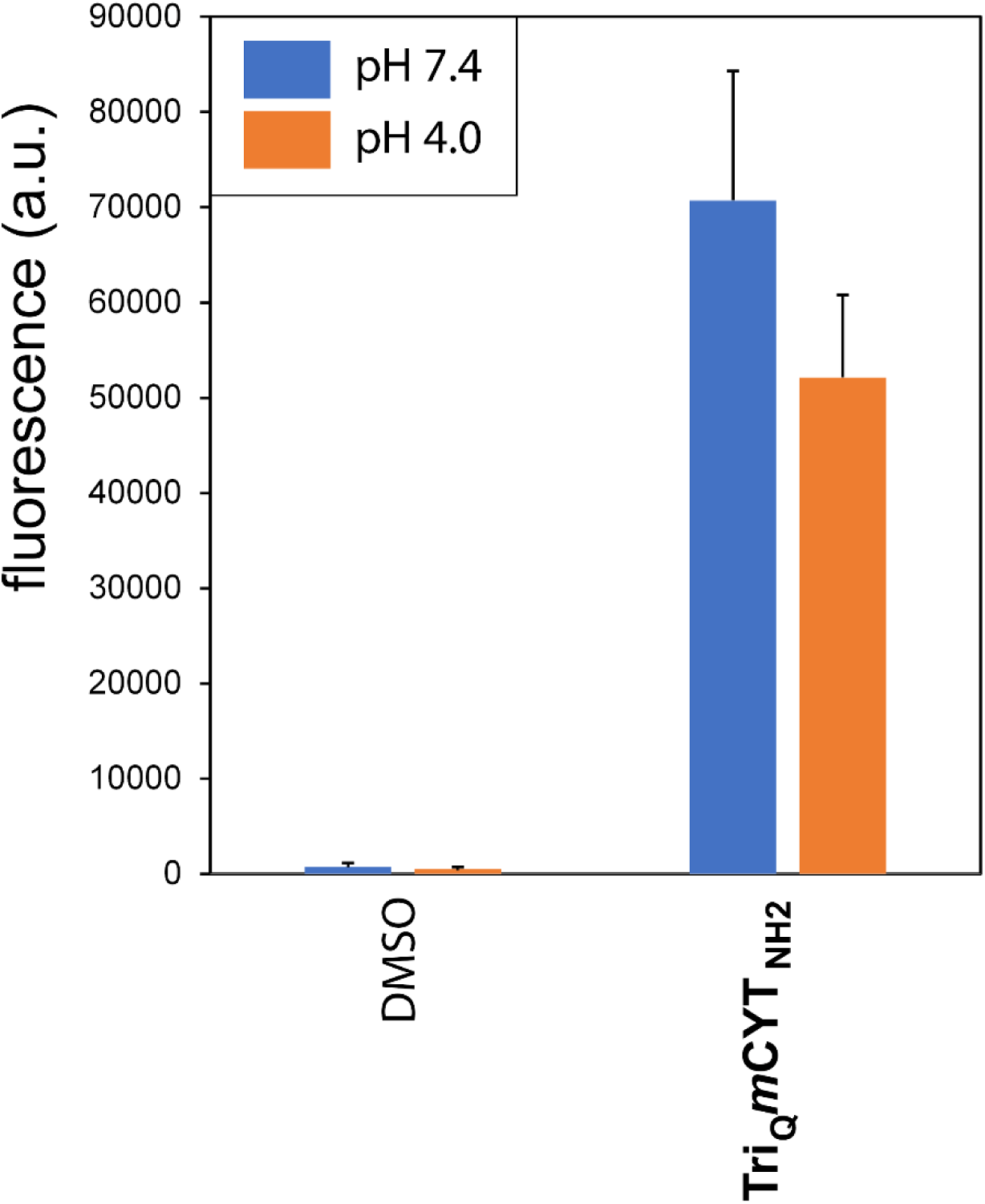
Flow cytometry analysis of *L. plantarum* treated overnight with 100 μM of **Tri_Q_*m*CYT_NH2_** at pH 7.4 or pH 4.0 (buffered with MES). Data are represented as mean + SD (n=3).

To confirm that the fluorescent signal in the flow cytometry experiments was, in fact, emitted from within the PG scaffold, a series of additional experiments were performed. First, *L. plantarum* cells were labeled with the tripeptide probe **Tri_Q_*m*CYT_NH2_**, fixed with formaldehyde to preserve the basic cellular structure, and subjected to a treatment with mutanolysin, which is expected to selectively depolymerize the PG scaffold.^68^ We found that fluorescent signals were reduced in a time-dependent manner, consistent with embedment of the probes within the PG scaffold (**Figure 9**). As a control experiment, labeled cells were treated with lysozyme and no reduction in cellular fluorescence was observed, as *L. plantarum* is inherently resistant to lysozyme due to *O*-acetylation of GlnNAc.^68^ As an alternative to PG digestion, which releases the entire scaffold regardless of what the stem peptide structure is, we took advantage of the disulfide bond to delineate probe incorporation. *L. plantarum* cells were incubated with **Tri_Q_*m*CYT_NH2_** and subsequently treated with dithiothreitol (DTT), which is expected to reduce the disulfide bond in the crosslinked *m*-CYT thus releasing the part of the probe containing fluorescein from the remaining PG scaffold (**Figure 10**). The observed decrease in cellular fluorescence is consistent with the proposed mode of incorporation and further suggests that the tripeptide probe is primarily incorporated as an acyl-acceptor strand because acyl-donor strands would not result in probe release from the PG. Finally, PG incorporation was verified using standard PG analysis from cells treated with **Tri_Q_*m*CYT_NH2_**. Two additional bacterial species were tested for *m*-CYT labeling including *Listeria monocytogenes* (*L. monocytogenes*) and *Lactobacillus casei* (*L. casei*). *L. monocytogenes* possess an *m*-DAP type of PG structure while *L. casei* is a L-Lys type bacteria that typically contains a D-Asn crossbridge.^59^ It was observed that *L. monocytogenes* labeled in a manner consistent with other *m*-DAP-based organisms (**Figure 11**) but, as expected, *L. casei* showed little labeling with the probe displaying *m*-CYT (**Figure 12**).

**Figure 9.**
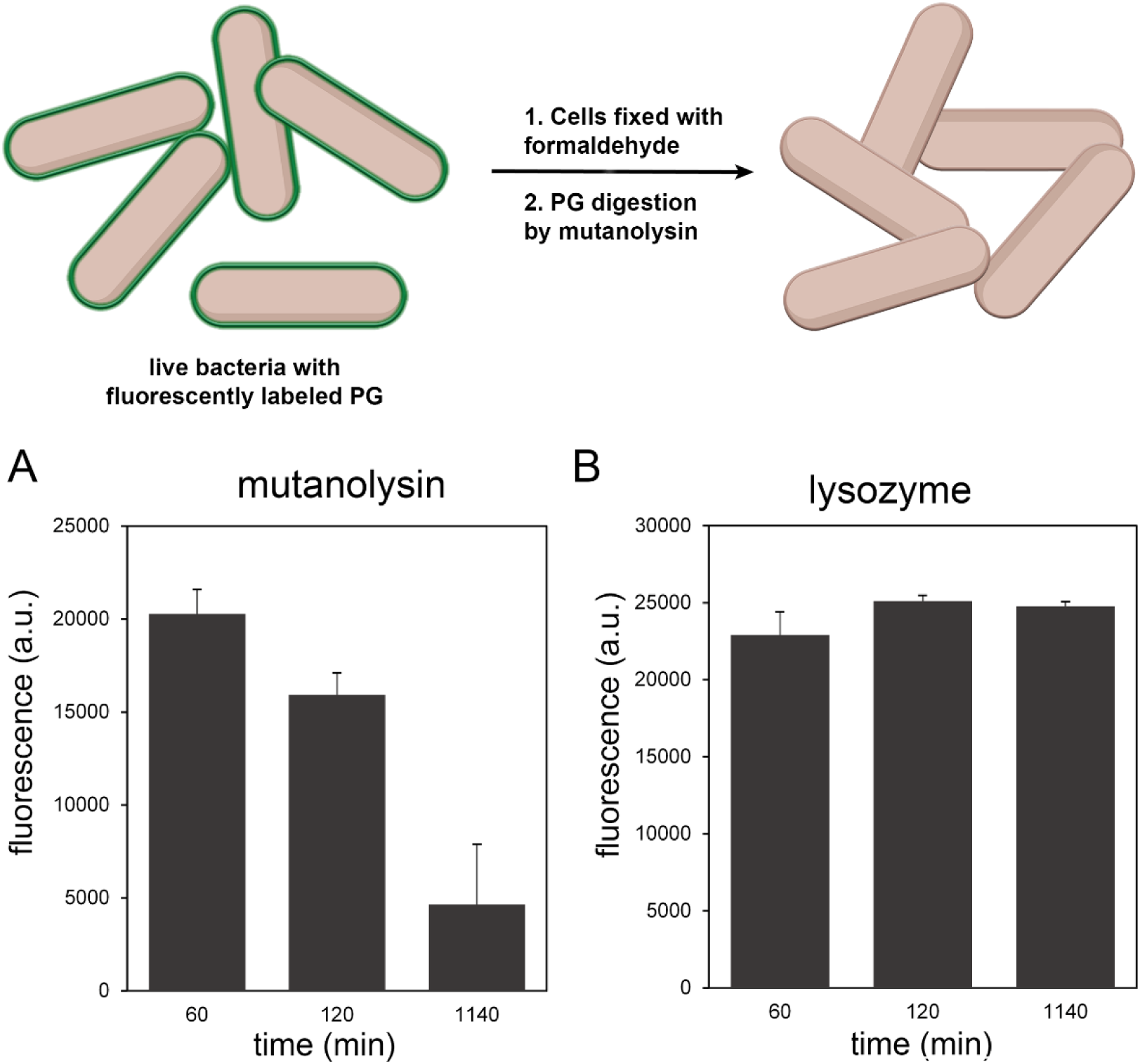
*Above*: Schematic of cells labeled and fixed with fluorescent probe, then treated with mutanolysin for PG digestion. (A) Flow cytometry analysis of *L. plantarum* cells labeled overnight with 100 μM **Tri_Q_*m*CYT_NH2_**, fixed with formaldehyde, washed 4X, and treated with 500 μg/mL lysozyme at 37°C. At various time points, samples were collected and cells were analyzed by flow cytometry. Data are represented as mean + SD (n=3). (B) Flow cytometry analysis of *L. plantarum* cells labeled overnight with 100 μM **Tri_Q_*m*CYT_NH2_**, fixed with formaldehyde, washed 4X, and treated with 500 μg/mL mutanolysin at 37°C. At various time points, samples were collected and cells were analyzed by flow cytometry. Data are represented as mean + SD (n=3).

**Figure 10.**
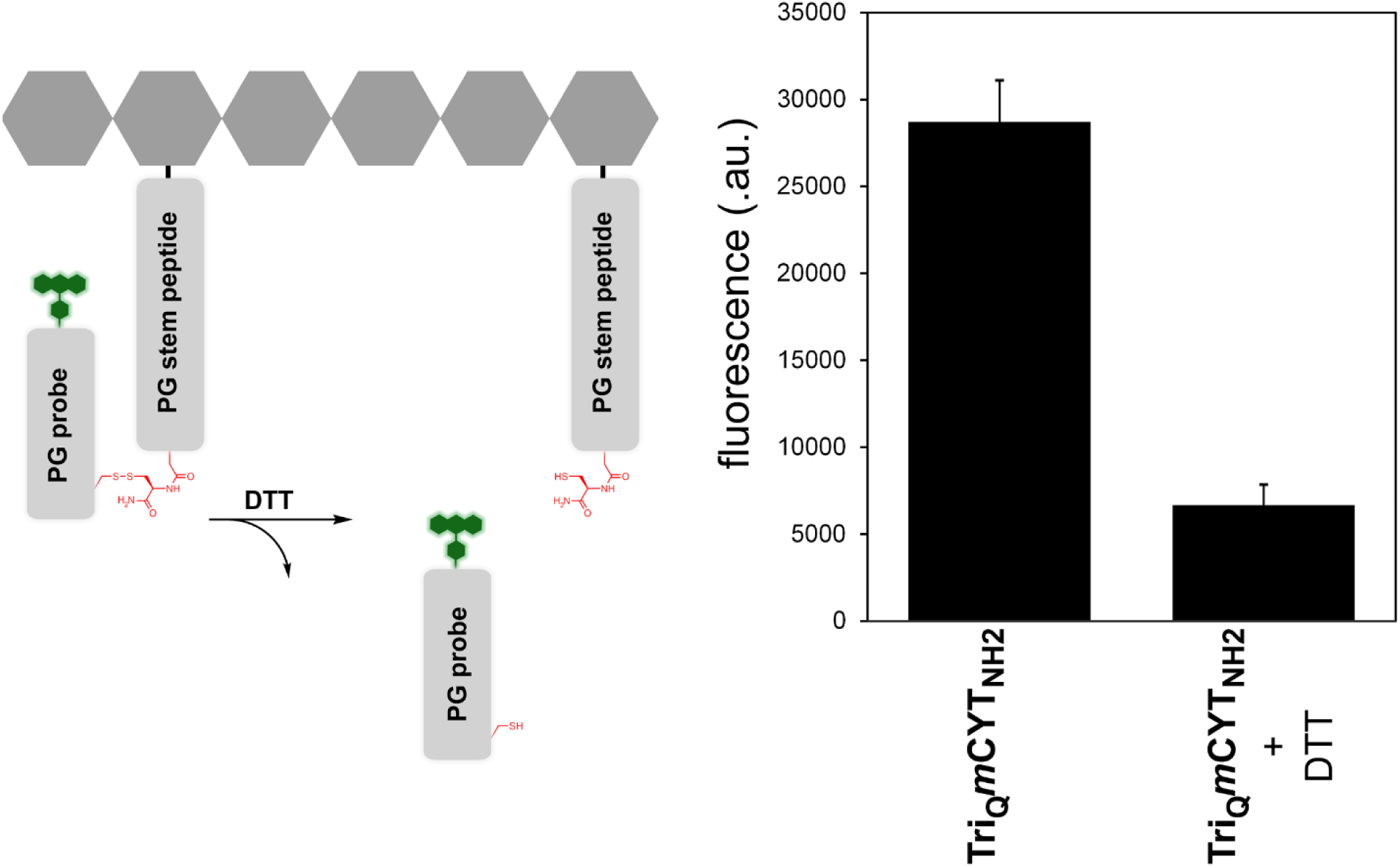
(*Left*) Schematic of **Tri_Q_*m*CYT** probe crosslinked to the PG stem peptide, followed by treatment of DTT which is expected to reduce the cystine and lead to release of the fluorescent probe. (*Right*) Flow cytometry analysis of *L. plantarum* treated overnight with 100 μM **Tri_Q_*m*CYT_NH2_**, fixed with formaldehyde and, if noted, treated with 10 mM DTT for 10 min. Data are represented as mean + SD (n=3).

**Figure 11.**
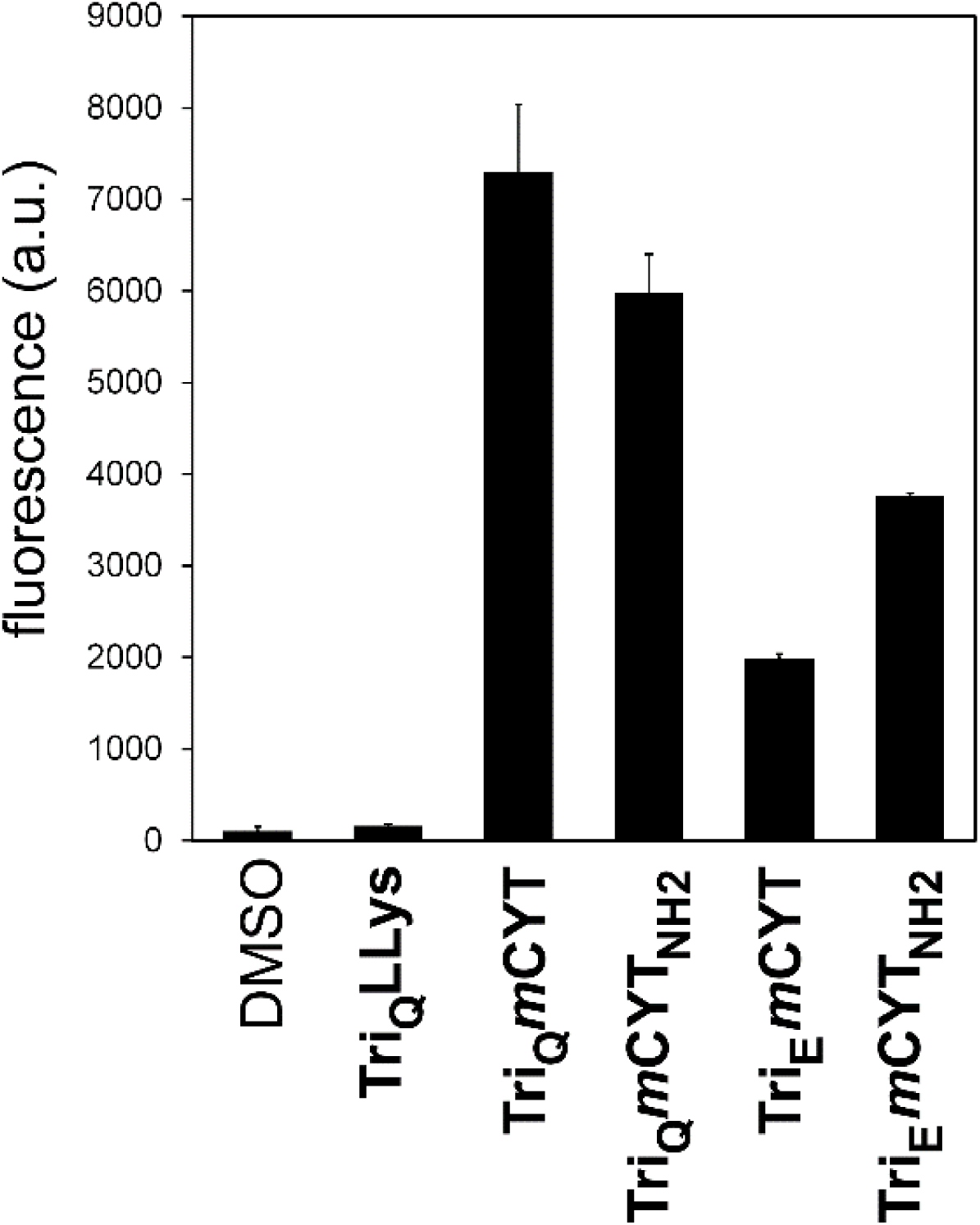
Flow cytometry analysis of *Listeria monocytogenes* treated overnight with 100 μM of each probe. Data are represented as mean + SD (n=3).

**Figure 12.**
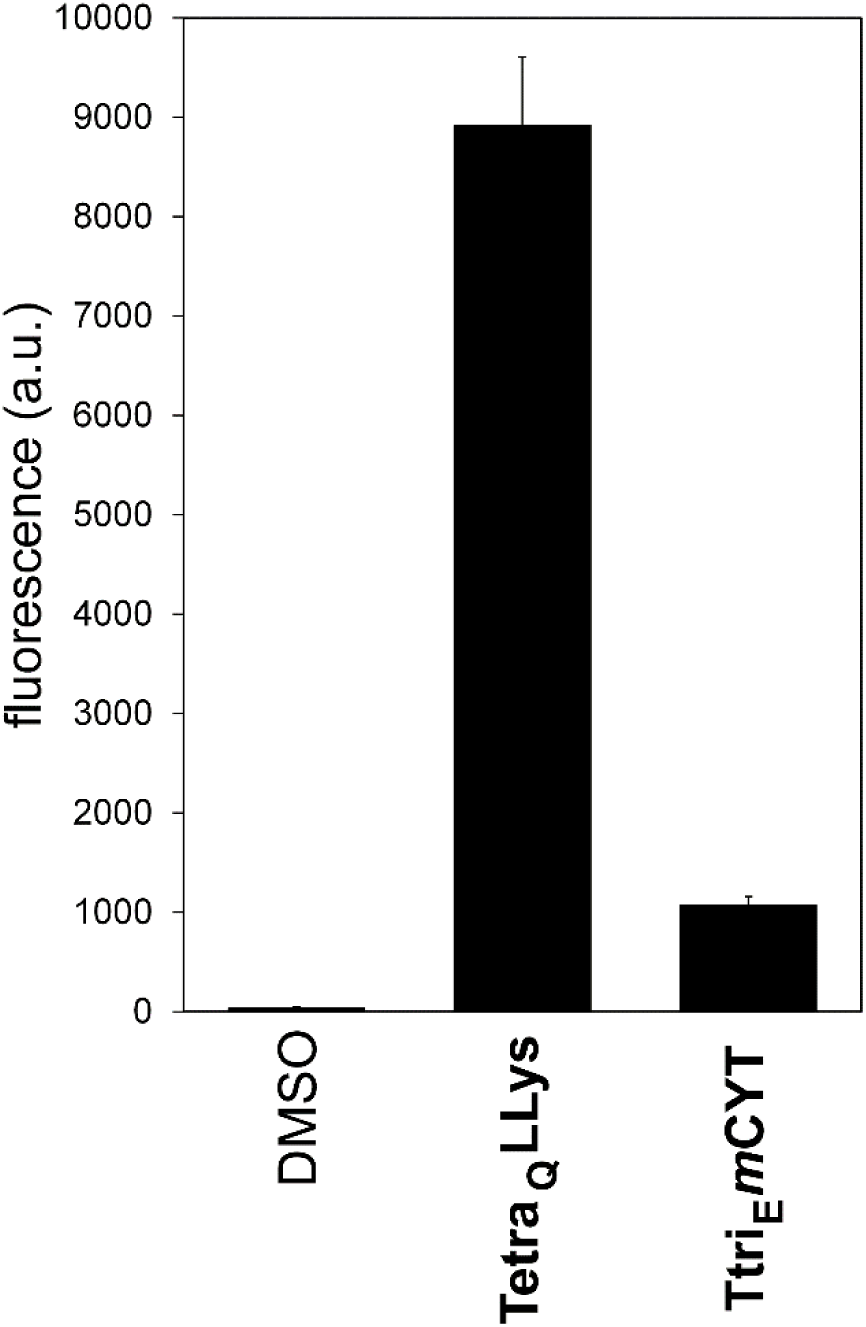
Flow cytometry analysis of *Lactobacillus casei* (*L. casei*) treated overnight with 100 μM of each probe. Data are represented as mean + SD (n=3).

There is increasing interest in understanding the cell wall biosynthetic machinery in live bacterial cells. The principal driving force for *in vivo* analysis is the emerging paradigm of extensive protein-protein interactions that control components of the cell wall biosynthetic machinery to modulate the growth and remodeling of PG. Our group and others have previously described a class of L-Lys-based PG probes that operates by mimicking components of the PG stem peptide to metabolically label the PG of live cells by TP-mediated incorporation, thus providing a direct signal that reports on PG biosynthesis. And yet, the lack of facile synthetic routes to *m*-DAP building blocks or their derivatives to assemble PG analogs of *m*-DAP containing organisms has hampered fundamental studies using live bacteria. We describe a strategy that makes use of the simple and widely available Fmoc-protected L-Cys to build *m*-CYT, which recapitulates key structural features of *m*-DAP. After demonstrating the ease of synthesizing *m*-CYT containing peptides, we showed that TPs accepted *m*-CYT in place of m-DAP by labeling live bacterial cells. More specifically, we installed *m*-CYT in a PG probe that can only be crosslinked into the bacterial PG scaffold by mimicking *m*-DAP. We anticipate that *m*-CYT will complement emerging tools to better understand cell wall biosynthesis in *m*-DAP containing organisms by providing a reliable route to a bioisostere that is biologically functional.

## ACKNOWLEDGEMENT

This study was supported by the NIH grant GM124893-01 (M.M.P.). Ldt-knockout strains were provided by Martin S. Pavelka (University of Rochester).

## FIGURES

**Scheme 1.**
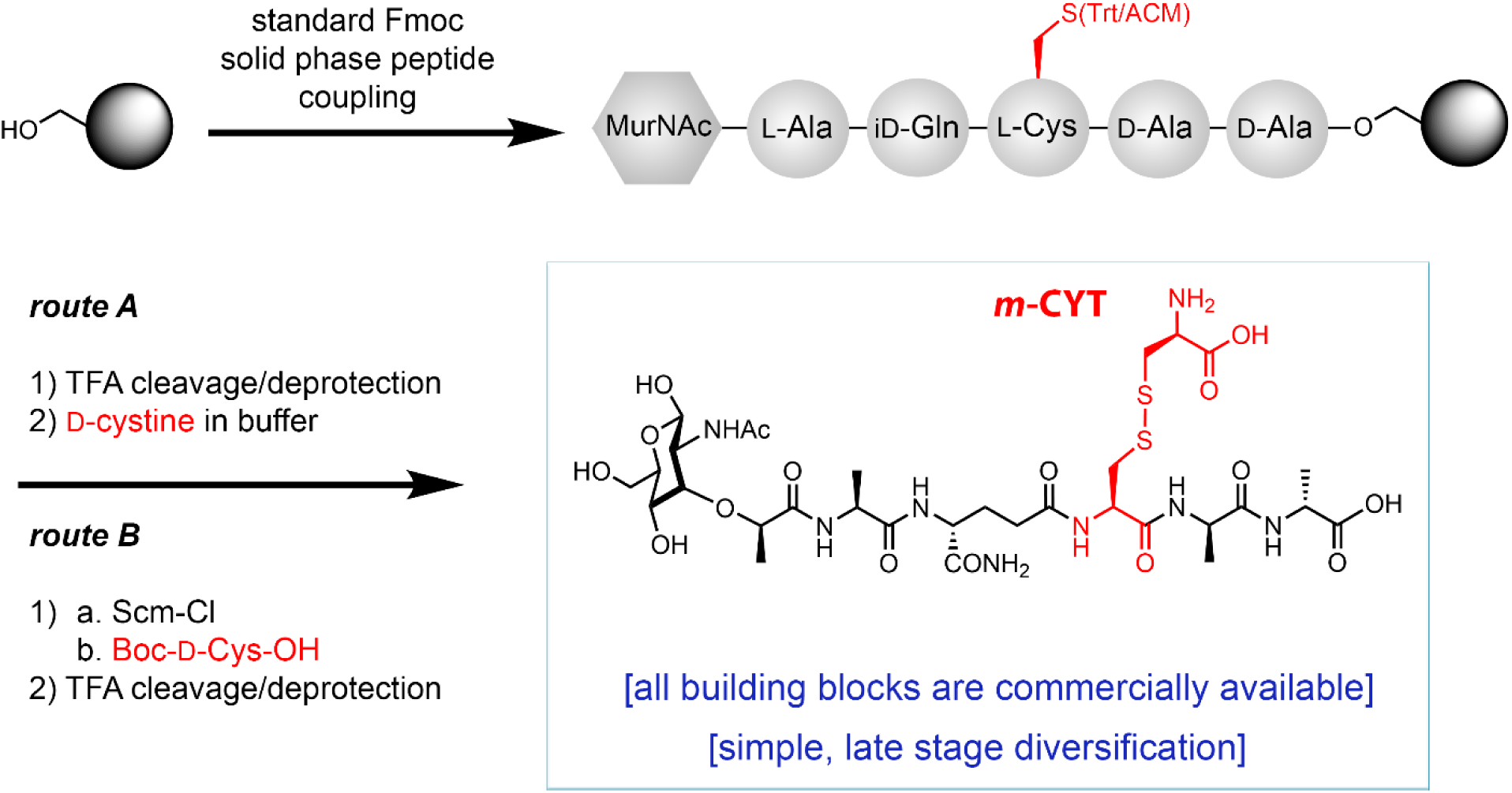
Two proposed synthetic routes to assemble *m*-CYT containing PG fragments.

**Scheme 2.**
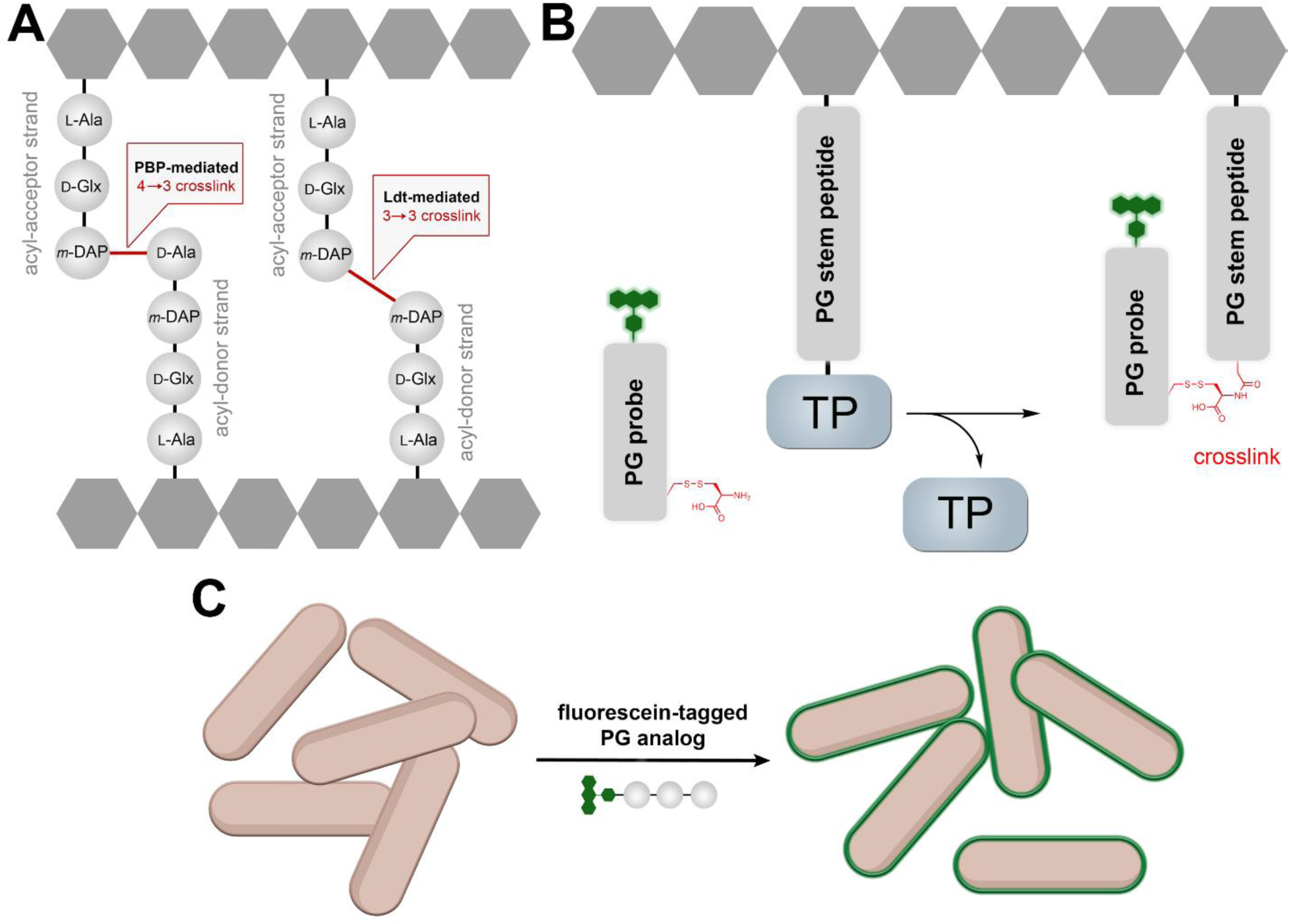
(A) Cartoon representation of the two primary modes of PG crosslinking carried about by D,D- and L,D-transpeptidases. (B) Snapshot representation of the acyl-intermediate of the TP reaction showing the hijacking of this step in the installation of the fluorescently tagged PG probe into the existing PG scaffold. (C) Probe incorporation on the cell surface can be readily measured using flow cytometry cell analysis.

